# Potential malaria vector *Anopheles minimus* (species A) still persisting in North East India

**DOI:** 10.1101/2020.09.01.277020

**Authors:** Varun Tyagi, Diganta Goswami, Sunil Dhiman, Dipanjan Dey, Bipul Rabha, P. Chattopadhyay, Sanjai K Dwivedi

**Affiliations:** Division of Pharmaceutical Technology, Defence Research Laboratory, Tezpur-784001, Assam, India; Medical Entomology Division, Defence Research Laboratory, Tezpur-784001, Assam, India; Vector Management Division, Defence Research & Development Establishment, Gwalior-474002, Madhya Pradesh; Cotton College State University, Panbazar, Guwahati-781001, Assam

**Keywords:** *Anopheles minimus*, species specific PCR, North East India

## Abstract

**Background:** Vector borne infectious diseases affect two third of the world’s human population and cause mortality in millions each year. Malaria remains one of the major killers in the Indian sub-continent and transmitted uninterruptedly by many efficient vectors and their sibling species. In North East India (NE), *Anopheles minimus* has been recognized as an important vector which shares majority of malaria cases. This study primarily focuses on to recognize the presence and distribution of sibling species of *An. minimus* in certain endemic area of NE India.

**Methods:** *Anopheles* species were collected and identified using available morphological keys. The genomic DNA was extracted from the mosquito specimen and used to perform species specific PCR (ss PCR) for molecular identification of major malaria vector *An. minimus* sibling species

**Result:** Morphological identification suggested the presence of *An. minimus sl* in low density in the study area. The specimen of *An. minimus* subjected to ss PCR confirmed the prevalence of only one sibling species namely, *An. minimus* A in Sialmari and Chandubi.

**Conclusion:** Though in low density, but malaria vector *An. minimus* is still present in certain endemic areas of NE India. The ss PCR assay employed presently suggested that *An. minimus* sibling species A is prevailing in the region. Presently used ss PCR assay was simpler, faster, cheaper and more readily interpreted than earlier assays. This information could be useful in understanding of current prevalence and distribution of *An. minimus* sibling species complex in NE region of India.

## Background

Despite comprehensive interventions, malaria cases have shown increase globally during the year 2017 (219 million; 95% confidence interval: 203-262 million) as compared to 2016 (216 million; 95% confidence interval: 196-263 million) and 2015 (211 million; 95% CI: 192–257 million) (WHO, 2018). Malaria attributable deaths largely remained unchanged to 4, 35,000 in 2017 as compared to 4, 51, 000 in 2016. India continue to contribute significant share to malaria episodes in South East Asia Region, and reported an about 0.85 millions of confirmed malaria cases in 2017. *Plasmodium falciparum* dominates the malaria transmission in India and accounts for >60% of malaria cases followed by *P. vivax* (>35% cases) annually. These two malaria parasites are uninterruptedly transmitted by six major vectors namely, *An. culicifacies, An. fluviatilis, An. stephensi, An. minimus, An. dirus and An. annularis* in different parts of the country (WHO, 2018).

North East region of India (latitude-21°58’ N to 29°30’N and longitude-88°3’ E to 97°30’ E) shares international border with many endemic countries, such as, Bhutan, China, Myanmar and Bangladesh that report considerable malaria cases every year. The region predominantly has humid sub-tropical climate and comprises of hills, wetlands, dense rain forests and forest fringes to support vector mosquito growth and proliferation through most of the year. Although the region inhabits 3.5% of country population but contributes 17.5% of total malaria deaths reported in India. Although several *Anopheles* species have been incriminated as malaria vector in the region during the recent years (Dhiman et al., 2012, 2016; Dev et al., 2010) and associated with the transmission of both *P. falciparum* and *P. vivax* in the endemic pockets, but historically *An. dirus* (monsoon species) and *An. minimus* (perennial species) have been regarded as important malaria vectors involved in majority of malaria infections. Both these vectors incriminated in different independent investigations unequivocally were major vectors in the region few years ago. However during the recent years the density of *An. minimus* and *An. dirus* has declined considerably. Studies have suggested that both these vectors have been either disappeared (Saxena et al., 2014; Dev et al., 2015) or prevailing in very low numbers which may not be sufficient to maintain perennial malaria transmission (Yadav et al., 2017; Dhiman et al., 2012). Furthermore, secondary malaria vectors which had insignificant epidemiological importance earlier were incriminated harboring malaria parasites and maintaining uninterrupted malaria transmission in the region (Dhiman et al., 2012, Yadav et al., 2017). Although many investigations (Dev et al., 2015; Yadav et al., 2017) could not collect *An. minimus* during their study, but its prevalence in low density below collectable limit mainly in ecologically suited forest fringed pockets cannot be overruled.

*An. minimus* sensu lato (Funestus group) comprises of *An. minimus* s.s (formerly known as *An. minimus* ‘A’), *An. harrisoni* (formerly *An. minimus* ‘C’) and *An. yaeyamaensis* (formerly *An. minimus* ‘E’) (Green et al., 1990; Harbach, 2004, Dutta et al., 2014). *An. minimus* ‘E’ is restricted to Ryukyu Archepelago of Japan and not involved in malaria transmission (Somboon et al., 2001), whereas species ‘A’ and ‘C’ are predominantly distributed throughout South East Asia region and responsible for malaria transmission. In North East region *An. minimus* ‘A’ has been incriminated as malaria vector and found associated in malaria transmission in many states of the region (Dutta et al., 2014).

Molecular identification of sibling species An alternative approach was adopted by Van Bortel et al. (2000) who used PCR amplification of the rDNA rDNA internal transcriber spacer 2 (ITS2) region followed by BsiZI restriction enzyme digestion to distinguish *An. aconitus*, *An. jeyporiensis, An. minimus A* and C*, An. pampanai, An. varuna* and subsequently *An. culicifacies* (Van Bortel et al., 2002).

Proper and accurate identification of any vector concerned with malaria transmission, its distribution, behavior, vector competency and relative abundance is essential for its successful management and control. However, these are not easy to explain owing to difficulties in morphologically distinguishing two species from one another and from the other closely related ones: *Anopheles aconitus*, *Anopheles jeyporiensis, Anopheles culicifacies, Anopheles varuna*, and *Anopheles pampani*. For instance, in central Vietnam, members of both *An. dirus* and *An. minimus* species complexes were earlier considered primary vectors. However, (Van Bortel et al., 2001) revealed the fact that those mosquitoes which were identified as *An. minimus* earlier were actually *An. varuna* (a member of the Funestus Group that also includes the Minimus Complex). *Anopheles varuna*, being an extremely zoophilic species in the study region cannot be regarded as a vector. This incorrect identification led to the misuse of rare and valuable resources and thus creating hindrances in vector management strategies. In order to mitigate the difficulty of morphological identification, varieties of molecular techniques have been developed for distinguishing *An*. *minimus* s.l. and other closely related species (Green et al., 1990; Van Bortel et al., 1999, 2000; Phuc et al., 2003).

The present study was aimed at investigating the presence of potential malaria vector *An. minimus* in the region and to identify its sibling species in northeastern region of India

## Methods

### Study area and collection of *Anopheles* mosquitoes

Current study was conducted during March 2016 to September 2016 (pre-monsoon and monsoon season). The adult *Anopheles* mosquitoes were collected from certain ecologically suited sentinel locations in Nameri (26.93° N - 92.88° E), Sialmari (26.71° N – 93.09° E), Jongakholi (25.98° N - 91.24° E), Chandubi (25.88° N - 91.43° E) areas of Assam (Fig. 1; Table 1). These areas have vast forest land, scattered tea meadows and forest fringed inhabited areas with many streams, rivers and small irrigation canals with grassy margins, thereby providing conducive breeding habitat for *An. mini*mus as well as for other *Anopheles* mosquitoes.

**Table 1.** Morphological identified *Anopheles* species, collected in CDC Trap and hand catch during the study

| S. No | Location | *Anopheles* species | CDC trap collection (%) | $\chi^2$ (p) | Indoor resting collection (%) | $\chi^2$ (p) |
| --- | --- | --- | --- | --- | --- | --- |
| 1 | Nameri | *An. jeyporensis* | 3 (4.5) | 51.6<br>($<0.0001$);<br>df=6 | 1 (2.6) | 16.7 (0.01);<br>df=6 |
|  |  | *An. aconitus* | 2 (3.0) |  | 2 (5.1) |  |
|  |  | *An. varuna* | 4 (6.0) |  | 5 (12.8) |  |
|  |  | *An. vagus* | 24 (35.8) |  | 12 (30.9) |  |
|  |  | *An. annularis* | 18 (26.9) |  | 7 (17.9) |  |
|  |  | *An. philippinensis* | 10 (14.9) |  | 7 (17.9) |  |
|  |  | *An. crawfordi* | 6 (9.0) |  | 5 (12.8) |  |
| 2 | Sialmari | *An. jeyporensis* | 7 (13.2) | 25.3<br>(0.0001);<br>df=5 | 2 (5.4) | 28.2<br>($<0.0001$);<br>df=5 |
|  |  | *An. minimus/ An. varuna* | 4 (7.5) |  | 0 (0.0) |  |
|  |  | *An. aconitus* | 4 (7.5) |  | 4 (10.8) |  |
|  |  | *An. varuna* | 6 (11.3) |  | 6 (16.2) |  |
|  |  | *An. vagus* | 18 (34.0) |  | 11 (29.7) |  |
|  |  | *An. annularis* | 14 (26.4) |  | 14 (37.8) |  |
| 3 | Jongakholi | *An. aconitus* | 1 (1.8) | 96.3<br>($<0.0001$);<br>df=3 | 2 (5.4) | 43.6<br>($<0.0001$);<br>df=3 |
|  |  | *An. vagus* | 12 (21.8) |  | 7 (18.9) |  |
|  |  | *An. annularis* | 40 (72.7) |  | 24 (64.9) |  |
|  |  | *An. crawfordi* | 2 (3.6) |  | 4 (10.8) |  |
| 4 | Chandubi | *An. varuna* | 6 (14.3) | 10.5<br>(0.01);<br>df=3 | 5 (13.2) | 9.19 (0.03);<br>df=3 |
|  |  | *An. vagus* | 18 (42.9) |  | 16 (42.1) |  |
|  |  | *An. annularis* | 10 (23.8) |  | 9 (23.7) |  |
|  |  | *An. philippinensis* | 8 (19.0) |  | 8 (21.1) |  |

**Fig. 1.**
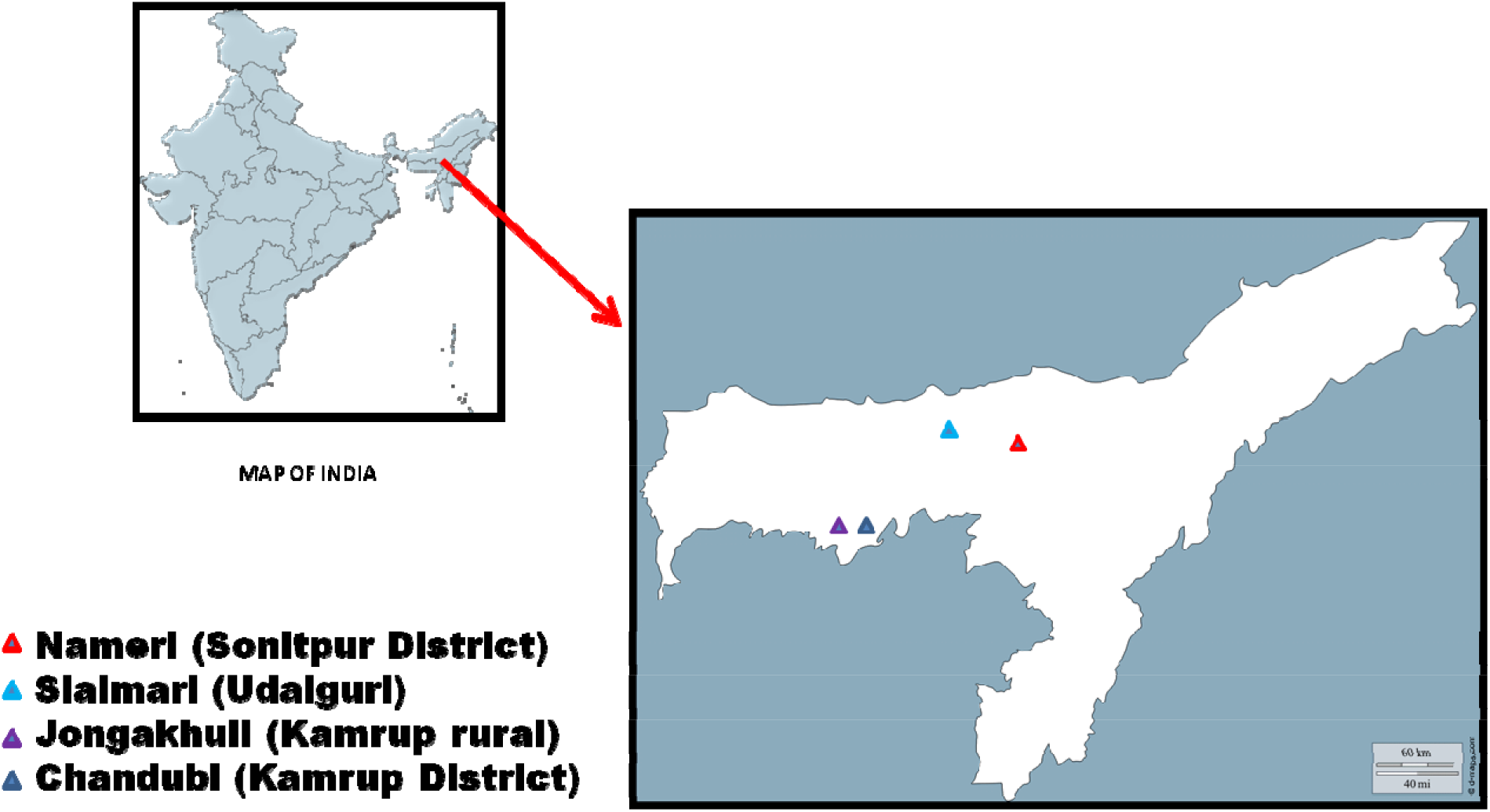
Collection sites: *Anopheles minimus* collection locations in Assam, North East India

The indoor resting mosquitoes were located by torch light and collected by using mouth aspirator during early morning (0500 – 0700 hr); whereas 6V battery operated CDC developed light traps were installed between 1800 hr to 0800 hr inside the human houses. The mosquitoes were morphologically identified using standard keys (Das et al., 1990; Harrison, 1980; Nagpal and Sharma, 1995). Since the study was primarily aimed at *An. minimus* mosquitoes, we separated mosquitoes that broadly look similar to *An. minimus*. These included *An. jeyporensis, An. minimus, An. acconitus and An. varuna* and broadly categorized into *An. minimus* group in the present study. These mosquitoes were first used to identify *An. minimus* s.l. and subsequently the identified *An. minimus* s.l specimens were processed for sibling species identification using species specific PCR assays.

### DNA Isolation

The DNA extraction was performed as described previously (Tyagi et al., 2016). Briefly, each adult female mosquito was homogenized by using polypropylene micro pestle (Tarson, India) in 2 ml micro centrifuge tube (Tarson, India) having filled with 100 μl lysis buffer containing 0.1 M Tris–HCl, 0.05 M EDTA, 0.2 M Sucrose, 0.05 % SDS, 0.1 M NaCl. The homogenate was immediately kept on ice for 10 min followed by heat treatment at 65 °C for 30 min. Subsequently, 30 μl 5M potassium acetate was added and immediately transferred to ice for one hour followed by centrifugation at 13,000 rpm for 15 minutes at 10 °C. A double volume of absolute chilled ethanol was added to the supernatant. The tube was left undisturbed for precipitation of DNA and stored at −20 °C for overnight. After centrifugation at 13,000 rpm for 15 min at 10 °C, the precipitated DNA was washed in 70% ethanol twice. The DNA pallet was allowed to air dry and finally dissolved in 50 μl TE buffer for use as DNA template in PCR assays.

### *An. minimus* species specific PCR assay

The assay employed *An. minimus* species-specific reverse primers along with an universal forward primer (Phuc et al. 2003) derived from highly conserved 5.8S coding region (Table 3). The PCR reaction was performed in 25 μl reaction volume containing: 1x PCR buffer (100 mM Tris-HCl (pH 8.3), 500 mM KCl), 0.2 mM NTPs, 1.5 mM MgCl_2_, 25 ng each of six primers, 0.625 U Taq, and 20 ng of DNA template (Phuc et al. 2003). The thermal cycle profile was optimized to the following conditions; 94 °C for 5 min; then 32 cycles of 94 °C for 1 min, 60 °C for 2 min 72 °C for 2 min and a final extension at 72 °C for 7 min. Ten of PCR products mixed with 2 μl of ethidium bromide were run on 1.0 % agarose gel and the results were visualized under UV–VIS gel documentation system (Syngene, G-Box, UK). The method by Phuc et al. (2003) is able to identify *An. minimus* s.s. (species ‘A’) and species ‘C’ of *An. minimus* complex and two other members of the *An. minimus* group (*An. aconitus*, *An. varuna*) along with *An. jeyporiensis*. However we have used primers that were specific to *An. minimus* species ‘A’ and ‘C’ only to identify species ‘A’, species ‘C’ or hybrid of ‘A’ and ‘C’ from the collected samples.

**Table 2.**
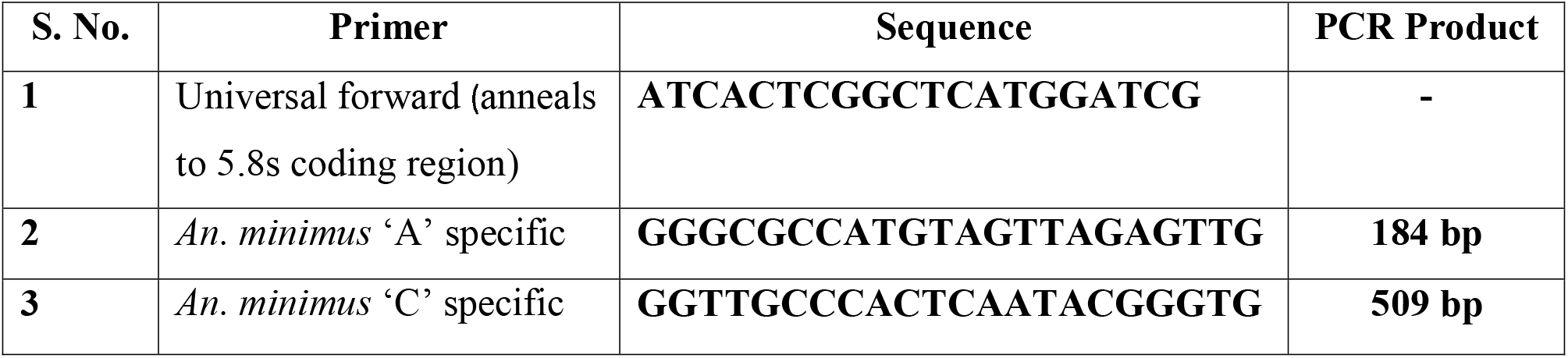
Detail of primers used for differentiating *An. minimus* ‘A’ and ‘C’ within *An. minimus* sibling species complex

## Results

### Mosquito collection

In the present study, we could collect specimens of different *Anopheles* species from the sentinel study sites using mouth aspirator and CDC light traps. Among the collected mosquitoes, two mosquito species namely, *An. annularis* and *An. vagus* were predominant and recorded in high number in all the collection points.

During the study, very few *An. minimus* mosquitoes could be collected. In Sialmari collection area, we found 4 *Anopheles* mosquitoes in bad condition and the morphological features of these specimens were deteriorated. Therefore we could not identify them correctly, however these were identified to *An. minimus* species ‘A’ using species specific PCR assay.

### Identification of *Anopheles minimus* sibling species

Species specific PCR for *Anopheles minimus* sibling species diagnosis used genomic DNA of field collected *An. minimus* s.l and amplified separately with primer specific for species ‘A’, species ‘C’ and both ‘A’ and ‘C’ for hybrid detection along with 5.8S forward primer for each reaction. First reaction with primers set ‘A’ yielded 184 base pair band that are specific to *An. minimus* species ‘A’ while PCR reactions for species ‘C’ and ‘A/C’ hybrid did not produce any band (Fig. 2).

**Fig 2.**
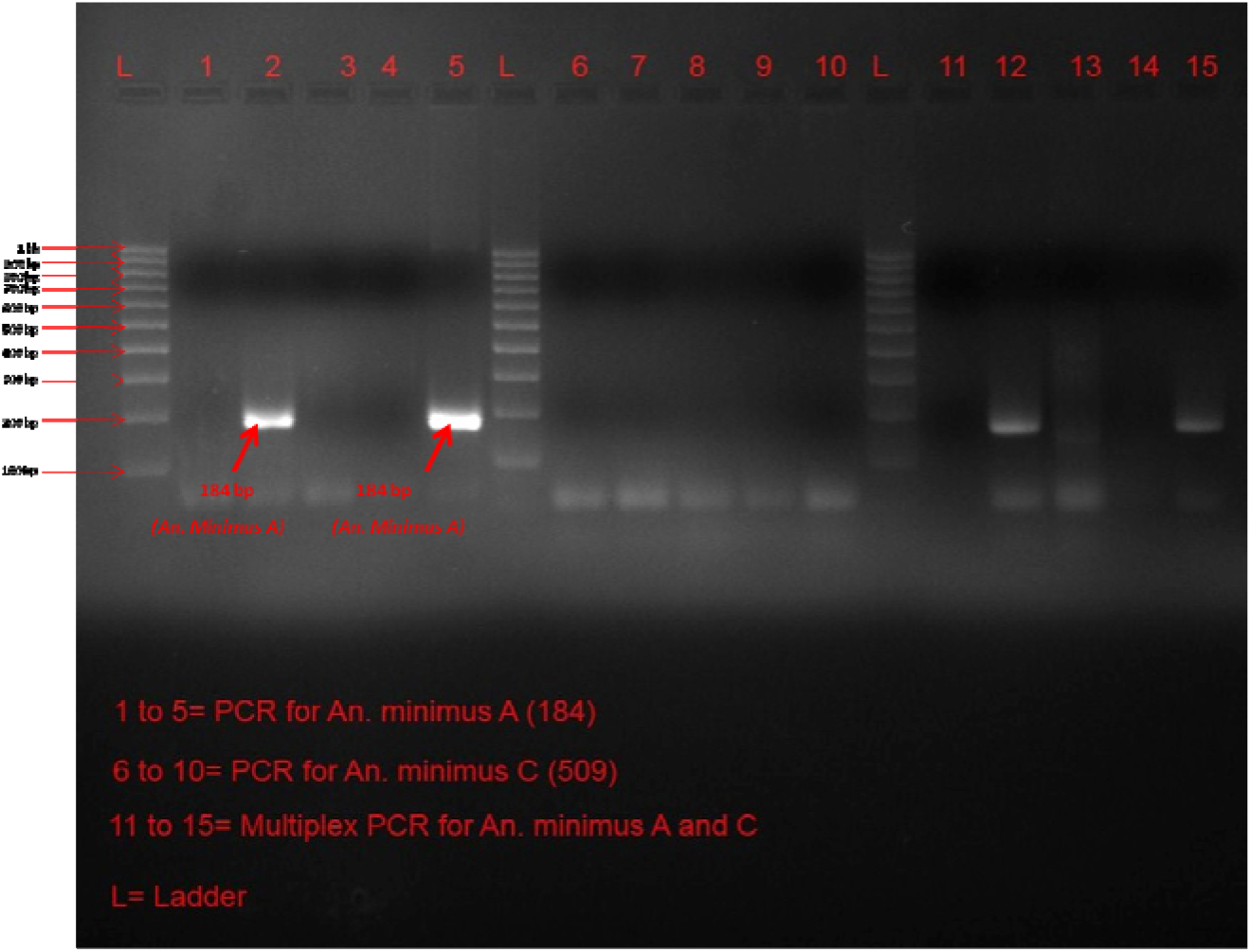
*An. minimus* sibling species identification using species specific PCR assay

## Discussion

*An. minimus* has been recognized as an efficient malaria vector in North East region of India since long and held responsible for perennial transmission of *Plasmodium* as evidenced by high sporozoite rates recorded in different seasons (Dev et al., 2016). In the region, *An. minimus* s.l. is endophillic and prefers to rest hiding on darker areas of walls, ceilings, cloths folding and temporary structures inside the human dwellings after taking blood meal. This species has very high affinity for human blood and has reported anthropophilic index of◻>◻90% in North East India (Dev et al. 1996, 2010; Dhiman et al., 2012). Considering this, the present collections intended to collect *An. minimus* mosquito were made inside the human houses. This species is still sensitive to insecticidal formulations used in intervention programmes; hence it is argued that *An. minimus* may be exhibiting behavioral changes by preferring to bite and rest outdoors than indoors in order to avoid direct contact with the indoor sprayed insecticides. Although investigations have reported significant declining trend in *An. minimus* density (Dhiman et al., 2012) even to virtually nil level in different states of North East region of India, however none of the study so far has evidenced that density has not declined but behavioral plasticity in *An. minimus* has led it to feed human blood and rest outdoor.

In the recent years there has been considerable change in the ecology in the region involving deforestation, increasing spread of cultivated lands and reduction in the breeding habitat, which has influenced the vector composition drastically (Nath et al., 2012; Yadav et al., 2012). *An. minimus* has high preference for egg laying places and lay eggs in grassy margins of unpolluted rivers and streams. Therefore water bodies lacking such supported ecology for *An. minimus* breeding may sometime account for absence of larvae in rivers and adults with in the flight range of this mosquito.

Species specific PCR assay used was able to identify species ‘A’ of *An. minimus* complex. Different PCR based methods have been used successfully in identification of mosquito complexes accurately (Walton et al., 1999; Chen et al., 2002; Phuc et al., 2003; Sharpe et al., 1999; Kengne et al., 2001; Singh et al., 2004; Tyagi et al., 2016). These methods are simple, precise and more accurately interpreted, hence cold be used in segregating the closely related sibling species from each other. Furthermore the multiplex PCR assay used presently was also able to identify sibling species ‘A’ of *An. minimus* among all the mosquito species used in the study. The multiplex PCR assay used here has produced unambiguous variation in PCR products of various *An. minimus* group mosquitoes in South East Asian countries, thus making this assay suitable and reliable for accurate identification of this mosquito complex (Phuc et al., 2003).

Studies have reported that *An. minimus* was once widely prevalent and involved in disease transmission in Himalayan foothills of northern to eastern region of India (Dev et al., 2016; Rao et al., 1984). However later thought to have been disappeared, as most of the studies did not report this species from Sub-Himalayan foothill regions (Devi et al., 2007; Mahesh et al., 2003; Shukla et al., 2007; Yadav et al., 2017). However, this species appeared in eastern region after more than four decades and involved in malaria transmission (Gunasekaran et al., 2014; Jambulingam et al., 2005). Similarly, malaria vectors that were disappeared completely and believed to be eliminated under the influence of control interventions or ecological changes were found to have been re-emerged later and incriminated as vector involved in malaria outbreaks (Hargreaves et al., 2000; Jambulingam et al., 2005).

Many studies conducted during past few years have reported malaria cases in North East region of India in the absence of *An. minimus*, while establishing that other anopheline mosquitoes that played insignificant role in malaria transmission previously, have taken over as important vectors and involved in continuous transmission of *Plasmodium* species throughout the year (Yadav et al., 2017; Dev et al., 2013, 2015). However the present study conducted in forest fringed foothills areas that provide suitable breeding and proliferation ecology for *An. minimus* s.l. confirms that this important vector has not disappeared completely but limited its prevalence into certain favourable areas. Present study has identified *An. minimus* species ‘A’ which is a well known vector in the region. The findings further suggest that *An. minimus* may be thriving under selection pressure as numbers of insecticidal products are in place to target this prominent vector. However it may re-surge in high density once these large scale interventions targeting this mosquito are withdrawn considering that it is disappeared completely.

## Conclusion

Present study has attempted to establish that *An. minimus* ‘A’ is existing in the North Eastern region of India but limited to certain ecologically suitable pockets in forest fringed areas. The environmental conditions still favour the prevalence of this potential vector which is probably striving under selection by changing ecology as well as increasing insecticidal pressure. Therefore complete disappearance of this mosquito in the study region can be overruled. Study also emphasize that it is not appropriate to conclude *An. minimus* group identification without using PCR like sensitive methods, as it may give confusing results uninvited for vector control programmes.

## Acknowledgements

We are thankful to the Director, Defence Research Laboratory Tezpur to support the work. The help rendered by the villagers during the study is deeply acknowledged.

## Authors’ contribution

VT: Developed study idea, performed laboratory experiments, prepared manuscript. DG: Developed study idea, involved in field collection, performed the experiments. SD: Developed study idea, involved in field collection, analyzed data and drafted manuscript. DD: Laboratory experiments and data analysis. BR: Field collection, identification and laboratory experiments. PC: Development of study idea, drafting of manuscript. SKD: Overall guidance and manuscript editing.

